# Using thermal death time models to analyze cold stress resistance across *Drosophila* species

**DOI:** 10.64898/2026.02.06.704298

**Authors:** Clara Garfiel Byrge, Louison Le Duff, Hervé Colinet, Mads Kuhlmann Andersen, Johannes Overgaard

**Author notes:** Author for correspondence Mads Kuhlmann Andersen.

## Abstract

- Chill-susceptible insects such as *Drosophila* are vulnerable to progressive disruption of ion and water homeostasis during cold stress, and low temperature exposure is a key factor affecting their physiology and distribution. Comparative studies of cold tolerance traditionally use simple or single-condition assays for interspecific comparisons, but the emergence of thermal death time (TDT) models offers a comprehensive framework to assess cold tolerance across different stress intensities and durations.
- Here we construct TDT curves for six *Drosophila* species, spanning boreal to tropical habitats, using Lt_50_ estimates across a range of stressful low temperatures (Lt_50_ ranging from ∼ 20 min to 2 days). For all species, the TDT curves provided good fits to the log(Lt_50_) vs. temperature data (R^2^ = 0.87 – 0.99).
- TDT curves from all species had steep slopes demonstrating that cold injury rate has a high thermal sensitivity such that small changes in temperature have profound effects on survival duration. The interspecific similarity of TDT slopes indicates that a conserved physiological dysfunction underlies cold injury across species. Further, additive accumulation of cold-induced injury in split-dose experiments suggests that acute and moderate cold damage represent the same underlying physiological dysfunction occurring at different rates.
- The TDT curve intercepts (species-specific tolerance thresholds) differed markedly between boreal, temperate, and tropical species and correlated strongly with their habitat temperature. Data from the present study and meta-analysis of published data find that the inherent species cold tolerance decreases by ∼ 0.45 °C for each °C colder the winter environment of the species is. When also considering the cold acclimation cues in cold climates we argue the experienced level of cold stress intensity is similar across environments inhabited by the *Drosophila* genus. This suggests that cold tolerance is important in shaping the fundamental niche of both boreal and tropical species.
- Overall, the TDT analysis of *Drosophila* at low temperature provides a powerful and predictive tool for quantifying insect cold tolerance. This approach enables detailed cross-species comparisons that allows for both ecological and physiological inference. Thus, TDT curves offer relevant approximations of insect cold resistance that could help predict insect responses to climatic change.

## Introduction

The body temperature of insects, and therefore the rate of their biological processes, depends on ambient temperature, making them sensitive to temperature changes in their environment (Stevenson, 1985; Angilletta, 2009). As a result, interspecific adaptations to tolerate and perform within specific temperature regimes represent key determinants of insect species geographical distribution. Multiple studies have found a strong correlation between measures of species thermal tolerance and the temperature of their habitat (Deutsch et al., 2008; Sunday et al., 2011; Sunday et al., 2012) including studies within the genus *Drosophila* that includes species ranging from tropical to boreal environments (Kimura, 2004; Kellermann et al., 2012a). The association between thermal tolerance and distribution is particularly strong with respect to cold tolerance limits. This is possibly due to the reduced ability to mitigate cold stress with behavior when activity rates are low. Additionally, winter represents a prolonged period of cumulative stress where the combination of cold and limited energy reserves likely defines the survival limit of a species. Consequently exposure to stressful cold in the environment can have direct impacts on survival or subsequent performance (Chen and Walker, 1994; Denlinger and Lee, 2010; Lee, 2012; Andersen et al., 2015; Koštál et al., 2019; El-Saadi et al., 2025).

Insects use various strategies to cope with cold stress. The most cold tolerant species are able to survive extreme low temperature by either tolerating ice formation within their extracellular fluids (freeze tolerance) or by lowering their supercooling point (SCP) and thus avoiding ice formation to very low temperatures (freeze avoidance) (Zachariassen, 1985; Ramløv, 2000; Denlinger and Lee, 2010). Most insects, however, suffer from cold at temperatures considerably above their SCP, making them ‘chill susceptible’ (Bale, 1996; Sinclair et al., 2003). The central challenge for chill susceptible insects is therefore directly related to temperature effects on biological rates and not related to the ice-water transition that is central for adaptations in freeze tolerant and freeze avoidant species (Bale, 1996; Denlinger and Lee, 2010). There is considerable interspecific variation in the level of cold tolerance among chill susceptible species (Sinclair et al., 2003; Overgaard and MacMillan, 2017). The most chill tolerant species can survive temperatures approaching their SCP (often at low subzero temperatures) whereas the most chill susceptible species succumb well above their SCP and for many tropical but also temperate species this may occur well above 0°C (Bale, 1993; Sinclair et al., 2003).

The degree of cold tolerance (ability to cope with stress) or cold resistance (the ability to lower the thermal threshold for when the cold is stressful) in chill susceptible insects has historically been characterized through a variety of measurements. These include lower lethal temperatures (LTemperatures), lower lethal times (Ltimes), the critical thermal minimum (CT_min_), and cold coma recovery time (CCRT) (Andersen et al., 2015; Sinclair et al., 2015). Most of these assays represent relatively acute measures of cold tolerance (minutes to hours), but it is well established that the estimation of thermal tolerance depends not just on the intensity of the stress but also on the duration that the stress is applied (Chen and Walker, 1994; Rezende et al., 2014; Enriquez and Colinet, 2017). To account for the combined effect of intensity and duration, cold tolerance can also be characterized using thermal death time (TDT) models, which integrate survival across a wide range of stress intensities and durations (Jørgensen et al., 2021; Tarapacki et al., 2021). In TDT analyses, mortality (e.g. Lt_50_) is measured at a series of fixed temperatures that represent different stress intensities and subsequently log_10_-transformed Lt_50_ values are plotted against exposure temperature to generate linear TDT curves that describe the exponential relationship between temperature (stress intensity) and Lt_50_ (**Fig. 1**).

**Figure 1:**
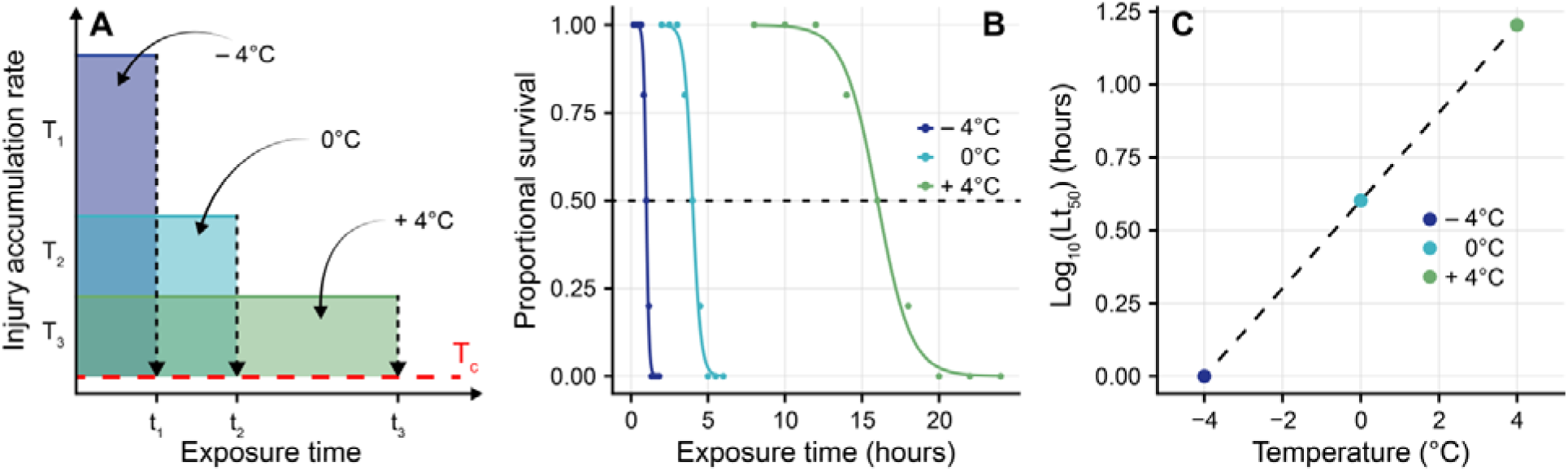
Construction of thermal death time (TDT) model for chill tolerance. A) Cold injury starts to accumulate once the ambient temperature drops below a species specific “critical temperature” (T_c_, red dashed line) after which injury accumulation rates increase exponentially with decreasing temperature. Intense cold stress (T_1_, purple square) can be tolerated briefly before the onset of injury and mortality (vertical dashed arrow, t_1_), while intermediate cold stress (T_2_, blue square) or even milder cold stress (T_3_, green square) can be tolerated for longer durations before mortality occurs (t_2_ and t_3_, respectively). Here the model assumption is that the total amount of damage tolerated (indicated by the area of the colored squares) is similar regardless of the stress intensity. B) Example data of Lt_50_ curves highlighting the relationship between exposure time and survival at three different cold stress intensities. C) TDT curve illustrating the linear relationship between exposure temperature and Log_10_(Lt_50_), constructed from the Lt_50_ values obtained in panel B.

TDT curves estimate two key descriptors that characterize thermal tolerance. Firstly, the intercept which parametrizes tolerance at a given duration, for example how cold should it be before lethal time is 1 hour for a specific species (intercept on x axis) or how long a specific species can survive a given temperature (intercept on y axis). Secondly, the slope (or its derivatives, the thermal sensitivity coefficient (z’) and the rate of change to a biological process with a 10°C increase in temperature (Q_10_’)) parametrize how rapidly cold-induced injury rates increase or decrease with changing temperature (Rezende et al., 2014; Jørgensen et al., 2021; Tarapacki et al., 2021). Consequently, well-fitted TDT curves can interpolate or extrapolate survival outcomes at untested exposure durations or temperatures if the model is not extended beyond reasonable limits (see discussion in Jørgensen et al 2021). Thus, TDT models offer a simple mathematical approach to study species-specific thermal limits under ecologically relevant conditions that is increasingly being considered also under fluctuating conditions (Ørsted et al., 2024; Arnold et al., 2025). The TDT approach can be used to conduct thermal risk assessments by calculating daily thermal injury potential based on regional and seasonal temperature variations in areas where local temperatures are known. This method, previously applied to aphids (Li et al., 2023), can be very useful for assessing how current and future climates may influence winter survival in various species. However, TDT models are time consuming to establish and require a lot of animals, and especially at low temperature, there is still limited information on how low-temperature TDT parameters (slope/intercept) differ among species adapted to different habitats and temperature preferences.

There is substantial evidence that cold damage in chill susceptible insects is linked to loss of ion-and water-balance that causes severe hyperkalemia and loss of membrane polarization (Koštál et al., 2004; Overgaard and MacMillan, 2017; Overgaard et al., 2021). Ionoregulatory failure is observed both at the organismal level (Koštál et al., 2004; MacMillan et al., 2015a; MacMillan et al., 2015b) and within the central nervous system (Armstrong et al., 2012; Robertson et al., 2023), and the cause of this homeostatic collapse has been linked to different physiological processes that are termed either “direct” or “indirect chill injury”. Briefly, direct chill injury is proposed to occur rapidly due to acute cold stress as a result of structural damage to membrane, cytoskeleton or protein integrity (Quinn, 1985; Hazel, 1995; Koštál, 2010) and it is suggested that such structural damage subsequently compromises ion balance (Overgaard and MacMillan, 2017; El-Saadi et al., 2023). In contrast, indirect chill injury, that occurs due to chronic cold stress, is considered to occur when there is a mismatch in active and passive ion transport rates such that there is a progressive loss of homeostasis that develops under moderate but long cold stress (Koštál et al., 2004; Koštál et al., 2006; MacMillan and Sinclair, 2011; Overgaard et al., 2021). It remains unclear whether intense acute stress (direct injury) and moderate chronic cold stress (indirect chill injury) are distinct processes or whether they represent the same processes that simply proceed rapidly or slowly depending on stress intensity. Here we propose that analysis of TDT data may offer indirect insights into this problem. Because the exponential nature of the TDT curve is analogous to an Arrhenius plot, we argue that distinct differences in the proximal physiological causes of stress are likely to manifest as “breakpoints” on the TDT curve (Jørgensen et al., 2021; Ørsted et al., 2022). Further, sequential exposure to intense and moderate cold stress (or *vice versa*) is likely to be additive if the stress is caused by similar physiological processes while distinct physiological causes are less likely to be additive.

In the present study we measure Lt_50_ across a wide range of cold stress intensities in six *Drosophila* species to establish species-specific TDT curves. The stress intensities applied are designed to cause mortality (Lt_50_) after durations ranging from minutes to days. The natural distribution of the six species span a broad range of thermal niches (boreal, temperate and tropical) and the species are chosen to represent both of the major subgenera *Drosophila* and *Sophophora*. Using the TDT data we explore patterns of cold resistance (intercepts of the TDT curves) and thermal sensitivity of cold tolerance (slope of the TDT curves) among and within species with the following hypotheses: Firstly; we hypothesize that there is a difference in intercept between species TDT curves which is supportive of fundamental difference in basal cold tolerance such that chill tolerant species may experience the same type of cold stress, but only when exposed to a much lower temperature. Secondly; we hypothesize that species have similar slopes of their TDT curves which is supportive of similar physiological cause of chill injury once the species experience similar levels of cold stress. Thirdly; we hypothesize that TDT curves are generally linear and uniform without obvious breakpoints. A uniform pattern supports the notion that cold stress (direct or indirect) is of similar nature across a range of stressful temperatures and that it is simply the rate of injury (homeostatic disruption) that differs. To support/falsify this latter hypothesis we also subjected flies to sequential exposures of intense and moderate cold stress to examine if the effects were additive (supportive of similar proximal causes).

## Materials and Methods

### Species characteristics and animal husbandry

Six species of *Drosophila* were used in this study (**Table 1**), ranging from the chill tolerant and boreal *Drosophila montana* and *Drosophila obscura* through temperate *Drosophila hydei* and *Drosophila melanogaster* with intermediate chill tolerance to the chill sensitive tropical species *Drosophila sulfurigaster* and *Drosophila teissieri*. Species were chosen such that both of the major subgenera *Drosophila* and *Sophophora* were represented for each “ecotype” and degree of cold tolerance (based on recordings of CT_min_ from Kellermann et al. (2012a) and 24h LT_50_ from Moos et al. (2025)). Data for average latitude, and average temperature of the coldest quarter for each species were obtained from Moos et al. (2025) that compiled geographical locations of the six species from published records sampled in the TaxoDros database (https://www.taxodros.uzh.ch/). These locations were then associated with climate variables using the WorldClim database (http://www.worldclim.org) (see details in Moos et al. (2025)). The same approach was used to obtain climate variables for species across different studies where these were not previously obtained.

**Table 1.** - Information on *Drosophila* species source populations, cold tolerance, natural latitudinal distribution, and natural environmental cold. ‘D.’ denotes genus *Drosophila*, whilst ‘(Dros)’ and ‘(Soph)’ indicate the two main subgenera *Drosophila* and *Sophophora*. Cold tolerance values for thermal critical minimum (CT_min_, the temperature at which the organism enters a comatose state) are sourced from Kellermann et al. (2012a), and the lethal temperature (LT_50_, the temperature leading to 50% mortality following 24 hours exposure), are obtained from Moos et al. (2025). Estimates of average latitude and average temperature of the coldest quarter for each species are sourced from Moos et al. (2025).

| Species (subgenus) | Country of origin | Laboratory/provider | CT<sub>min</sub> (°C) | LT<sub>50</sub> (°C) | Avg. latitude (°) | Avg. temp of coldest quarter (°C) |
| --- | --- | --- | --- | --- | --- | --- |
| *D. montana* (Dros) | Norway/Finland | Kankare/Hoikkala | -3.06 | -8.98 | 47.43 | -4.98 |
| *D. obscura* (Soph) | Finland | Longdon/Lankinen | -0.47 | -8.60 | 50.52 | -0.04 |
| *D. hydei* (Dros) | Japan | Kristensen | 2.15 | -2.98 | 36.55 | 6.19 |
| *D. melanogaster* (Soph) | Denmark | Sørensen/Overgaard | 1.91 | 0.06 | 35.54 | 6.24 |
| *D. sulfurigaster* (Dros) | Australia | Hrček | 6.84 | 6.99 | 17.10 | 18.84 |
| *D. teissieri* (Soph) | N.A. | Longdon | 6.94 | 3.94 | 6.29 | 21.69 |

All species were maintained under common garden conditions at 19°C under a 22:2 h light:dark cycle. Parental flies were reared in 250 mL plastic bottles containing 70 mL Leeds medium (recipe: 1 L water, 60 g yeast, 40 g sucrose, 30 g oatmeal, 16 g agar, 12 mL nipagin, and 1.2 mL acetic acid; Moos et al. (2025)) and these flies were transferred onto new food every seven days. For experiments, only female flies were used; newly hatched flies were collected every 2-3 days post emergence and briefly anesthetized with CO_2_ (< 10 minutes) during which they were sorted by sex under a microscope. Flies were divided into batches of ∼7-12 individuals and placed into 25 mL plastic vials containing 5 mL Leeds medium. They were then maintained at 19 °C for at least two days to recover from the anesthesia and handling and all flies were used for experiments at the age of 6-10 days post emergence. Note, however, that flies used for examination of longevity at different temperatures were placed directly at their designated temperature after anesthesia (see below).

### Assessment of Lt_50_ and construction of thermal death time curves

To construct thermal death time (TDT) curves for cold stress, we first assessed Lt_50_ across a range of temperatures for each species using an approach similar to Tarapacki et al. (2021). Briefly, at each experimental temperature 6-22 vials with foam stoppers (each containing 7-12 female flies; see **Table S1** for more details) were submerged into a temperature-controlled bath filled with a water-and-ethylene-glycol mixture pre-set at the desired experimental temperature after which vials were removed at different time points (**Fig. 1B**). After exposure, flies were transferred back into food vials at 19 °C and placed horizontally to avoid flies getting stuck in the food. Approximately 24 hours after the cold treatment, mortality was assessed based on their righting ability: Flies able to stand were considered alive, while immobile or impaired flies were considered dead (Colinet et al., 2017). Each species was exposed to 6-8 different experimental temperatures spanning from intense to mild cold stress and at each temperature, exposure times were chosen to secure data points with high, intermediate, and low survival, ensuring coverage of the expected Lt_50_ values ranging from ∼20 minutes to ∼2 days. When the experimental temperature was set below 0°C, a piece of filter paper was added along the edge of the vial to reduce the risk of flies being exposed to ice crystals from condensed water on the vial wall as this could cause inoculative freezing (Lebenzon et al., 2023). At temperatures above CT_min_, a filter paper with a droplet of Leeds medium was added to prevent starvation or dehydration. Temperature was monitored using a Testo 108 thermometer placed into an empty vial in the temperature bath, and each reported experimental temperature represents the average of four measurements.

Lt_50_ was calculated in R software (version 4.5.2, (R Core Team, 2025)) using the nlsLM() function from the minpack.lm package (Elzhov et al., 2016) where proportional survival against time was fitted to a logistic regression model using the following function:

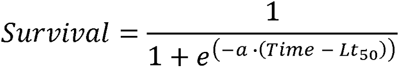

where *a* is the slope at the midpoint of the logistic regression and Lt_50_ is the time at which survival is 50%. This approach was used across all species and at each temperature tested. In four cases, however, the model showed standard errors orders of magnitude larger than the predicted values for a and Lt_50_ (*D. melanogaster* at −4.7°C and 4.0°C, *D. hydei* at −4.7°C, and *D. teissieri* at 5.8°C). This was solved by the removal of a single data point from each survival curve during the analysis (11 h and 148 h for *D. melanogaster*, respectively, 167 h for *D. hydei*, and 24 h for *D. teissieri*) which had negligible effects on mean values but reduced standard errors to a fraction of the means.

Subsequently, for each logistic regression we calculated a variance-based R^2^ as follows, in order to estimate the goodness of each logistic model:

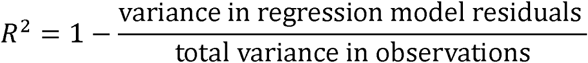

To construct the final TDT curves, the Lt_50_ values were log_10_-transformed and plotted against the exposure temperature for each species after which the relationship was described with linear regression (example data in **Fig. 1C**). Subsequently, we used the TDT regression models to calculate the thermal sensitivity, z’, for cold tolerance (Rezende et al., 2014; Tarapacki et al., 2021) which we subsequently converted into Q_10_’ as follows:

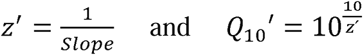

The’ signifies that the rate of chill injury accumulation increases when temperature decrease and the Q_10_’ represents the factorial change in an injury rate per 10 °C *decrease* in temperature (Rezende et al., 2014; Jørgensen et al., 2021).

The fitted TDT curves allow for estimation of stressful temperatures that would cause 50% mortality after fixed and mixed durations of exposure for each species (e.g. 30 or 300 min), to assess the time a fixed temperature (e.g. −6 °C) could be tolerated or the temperature needed to cause LT_50_ after a specific duration (e.g. 12 hours) (see analysis below)

### Assessment of cold damage additivity

To investigate if cold injury sustained at different cold stress intensities is additive, we exposed flies to a combined treatment with the hypothesis that flies receiving half of the Lt_50_-dose (i.e. time to 25% mortality) at two different and subsequent cold stress intensities would also lead to 50% mortality (Lt_50_). Specifically, we used the species-specific TDT curves to calculate the experimental temperatures corresponding to an Lt_50_ of ∼30 minutes and ∼300 minutes respectively (**Table S2**). We then exposed replicates of flies (8-12 females per replicate, 6-8 replicates per treatment) to one of five treatments: T_1_ = Intense cold stress for 30 minutes, T_2_ = Moderate cold stress for 300 minutes, ½T_1_ + ½T_2_ = Intense cold stress for 15 minutes followed by moderate cold stress for 150 minutes, ½T_2_ + ½T_1_ = Reversed order of the previous treatment. Finally a control treatment (C) was included where flies were handled by transferring them between vials and leaving them at room temperature for 330 minutes without exposure to cold stress. After all treatments flies were left to recover for 24 hours before survival was determined as described above.

### Longevity

To demonstrate that the relation between survival time and temperature is different in the permissive range of temperatures where there is no chill injury accumulation, we performed a longevity experiment to assess Lt_50_ at 11°C, 15°C, 19°C and 25°C. For each temperature and for each species ∼40-50 newly hatched female flies were split and placed into two 25 mL plastic vials (∼ 20-25 in each), containing 5 mL Leeds medium. A piece of filter paper was added to absorb excess moisture and allow climbing. Vials were placed at one of the four experimental temperatures in temperature cabinets with a 22:2 h light:dark cycle and fly mortality was recorded 3 times per week until the last fly died. Mortality during the first four days was disregarded and assumed to be a result from anesthesia and/or handling. To avoid flies getting stuck in the food they were moved to fresh food vials every 2-3 days with the exception of flies at 11°C that were only given new food once per week. For each species at each experimental temperature, Lt_50_ was assumed to be the day mortality surpassed 50%, and log_10_(Lt_50_) was plotted against temperature to provide TDT-like curves for longevity at permissive temperatures.

### Data analysis

All data analyses were performed using R software (v 4.5.2). Assessment of main effects of species and temperature and interaction in the log_10_-transformed TDT curves was performed using a linear model. To test for differences among species in cold tolerance characteristics, the slopes of the log_10_-transformed TDT curves (thermal sensitivity) were compared pairwise using the emtrends() function from the “emmeans” package (Lenth, 2023). A similar analysis was performed for the longevity experiment, except slopes were not compared across species. To further compare the cold tolerance of species and correlate this to their thermal environment, we used the linear regression equations from the TDT curves to estimate LT_50_ and Lt_50_ values at times and temperatures where most species overlapped on their exposures (12 h for LT_50_ and −6°C for Lt_50_). Where standard errors could not be directly extracted from models or transformed (e.g. from the log_10_- to the normal scale) the delta method was used to generate standard error estimates (Greene, 2003). Subsequently we tested these values, along with the TDT slopes against the species-specific average temperature of the coldest quarter and average latitude using Spearman’s correlations. Cold damage additivity within each species was analyzed using one-way ANOVA, followed by Tukey’s Honest Significant Difference (HSD) *post hoc* tests to investigate if the five additivity treatments differed. The level for statistical significance was 0.05 for all analyses, and error estimates presented or depicted represent standard errors of the mean unless otherwise specified.

## Results

### Cold tolerance assessment

The six *Drosophila* species were exposed to a range of cold temperatures for varying durations after which mortality was assessed to estimate Lt_50_. For each species we report Lt_50_ estimates ranging from ∼20 minutes to ∼2 days, and we find decreasing Lt_50_ durations with decreasing temperature across all six species. Overall, the survival data were well fitted to the logistic regression model, presenting sigmoidal Lt_50_ curves (**Fig. 2**) with an average R^2^ of 0.87 (median = 0.96 and range = 0.18 – 1.00). Out of 37 logistic regressions across species and temperatures R^2^ values were above 0.80 in 30 cases (∼ 81%, see **Table S1**).

**Figure 2.**
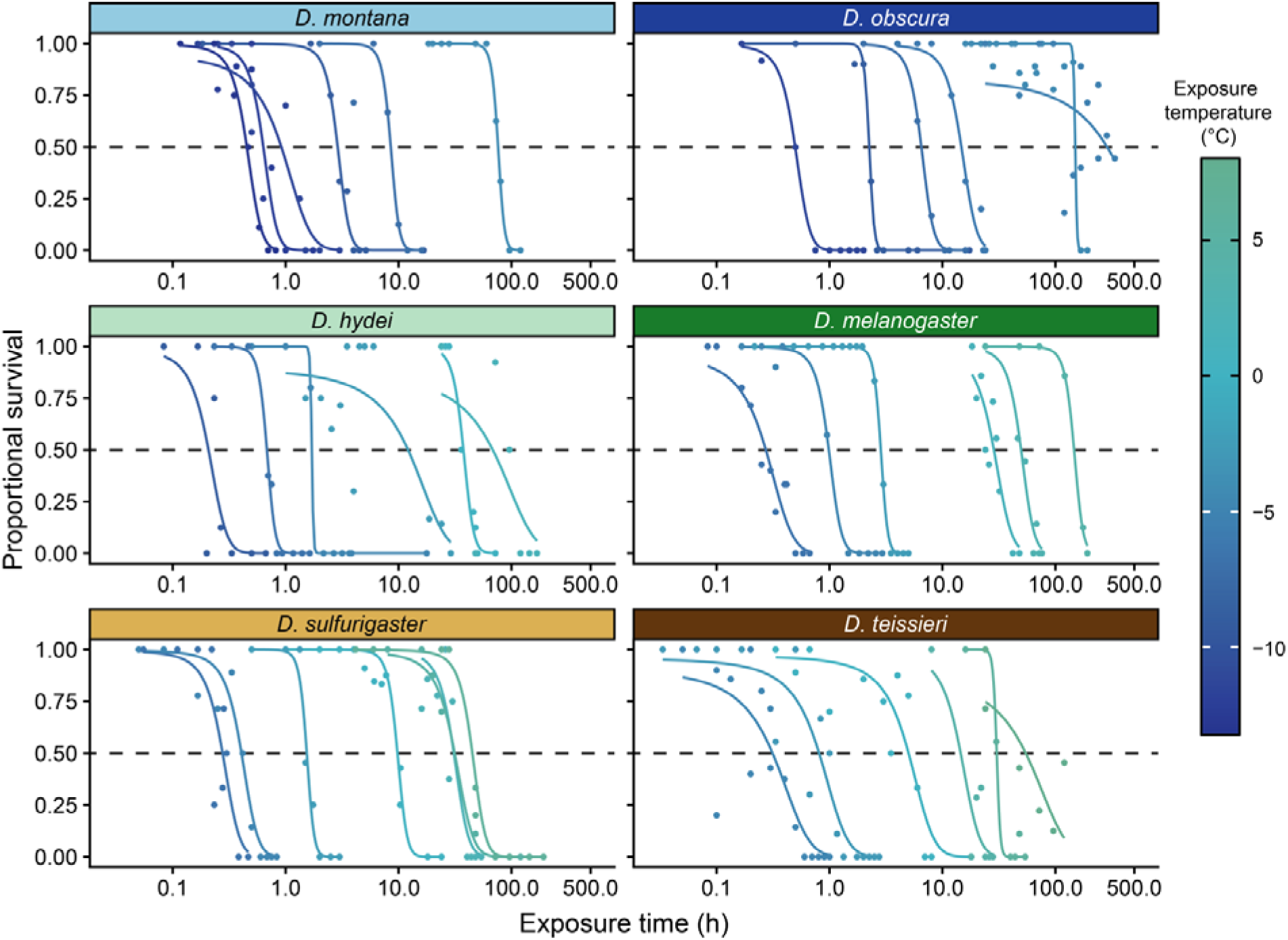
– Lethal time (Lt_50_) curves. Proportional survival plotted against exposure time at various temperatures for the six *Drosophila* species. Logistic regression models are fitted to the data points, and the time to 50% survival is indicated by the dashed horizontal lines. The average R^2^ was 0.87 (median = 0.96 and range = 0.18 to 1.00, based on 37 regressions).

### Thermal death time curves

The Lt_50_ estimates were log_10_-transformed and plotted against exposure temperature to generate TDT curves for each species (**Fig. 3**). All linear regression lines (TDT curves) were well-fitted to the log_10_(Lt_50_) vs. exposure temperature with R^2^ values ranging from 0.87 to 0.99 (**Table 2**) such that the TDT curves confirmed a positive, exponential relationship between Lt_50_ and exposure temperature (effect of temperature: F_1,25_ = 504.5, *P* < 0.001). There were clear differences between the TDT curves of the six species (effect of species: F_5,25_ = 5.7, *P* = 0.001) such that it required a lower temperature for the boreal species (*D. montana* and *D. obscura*) to experience a cold stress similar to that of the tropical species (*D. sulfurigaster* and *D. teissieri*) with the temperate species in between (*D. hydei* and *D. melanogaster*). We hypothesized that all six species would show the same thermal dependency of log(Lt_50_), however, our analysis showed that species differed (effect of species x temperature interaction: F_5,25_ = 5.5, *P* = 0.002). Pairwise comparisons of the TDT slopes revealed similar slopes across 5 of 6 species with the exception that the slope of *D. obscura* data was significantly steeper (higher thermal sensitivity) than that found for the two tropical species (see **Table 2**), though this is likely due to the unusually high Lt_50_ measured at −5.35°C.

**Figure 3.**
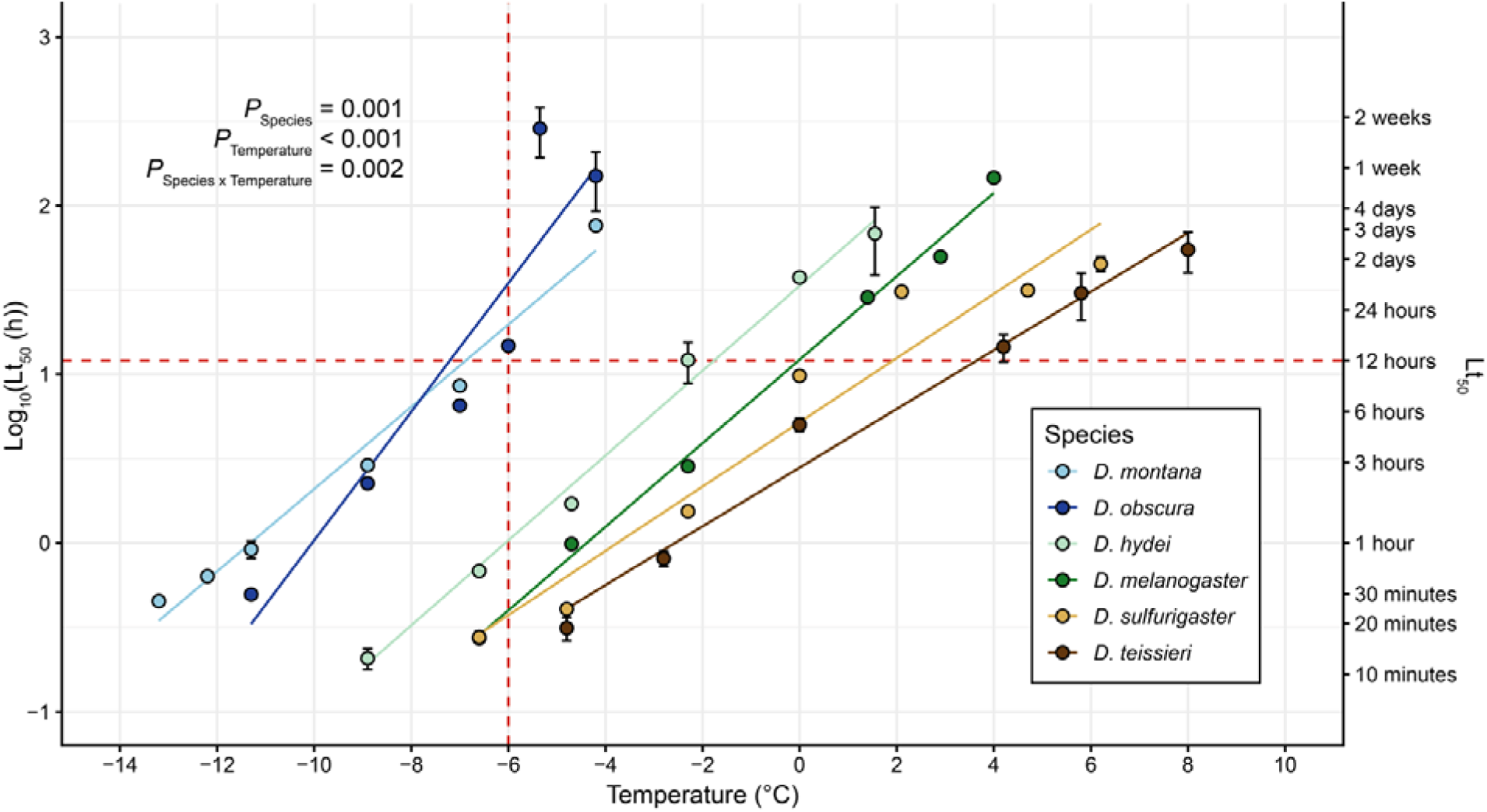
- Thermal death time (TDT) curves. Log_10_(Lt_50_) estimates (±SE) from the Lt_50_ curves (**Fig. 2**) plotted against exposure temperature for the six *Drosophila* species. Boreal species are shown in blue shades, temperate species in green and tropical species in brown. The intercept between the horizontal red dashed line and the TDT curves mark the species-specific LT_50_(12h), while the intercept between the vertical red dashed line and the TDT curves mark the species-specific Lt_50_(−6°C). Error bars not visible are obscured by the symbols.

**Table 2.** – Parameters of the thermal death time (TDT) curves for six *Drosophila* species. R^2^ values for the linear regression models, and species-specific values for the slope and two intercepts and their standard errors (horizontal = LT_50_ (12h) and vertical = Lt_50_ (−6°C)) presented in **Fig. 3**. The coefficients of thermal sensitivity, z’ and Q_10_’, are both derived from the slope. Shared superscript letters for the slope denote statistically similar values.

| Species | $R^2$ | Slope ($\log_{10}(\text{h})^{\circ}\text{C}^{-1}$) | $Lt_{50}(12\text{h}) (^{\circ}\text{C})$ | $Lt_{50}(-6^{\circ}\text{C}) (\text{h})^*$ | $z'^*$ | $Q_{10}'^*$ |
| --- | --- | --- | --- | --- | --- | --- |
| *D. montana* | 0.98 | $0.24 \pm 0.03^{ab}$ | $-6.93 \pm 0.28$ | $19.67 \pm 3.60$ | $4.10 \pm 0.50$ | $273 \pm 188$ |
| *D. obscura* | 0.87 | $0.38 \pm 0.04^a$ | $-7.20 \pm 0.43$ | $34.61 \pm 15.58$ | $2.62 \pm 0.27$ | $6454 \pm 5874$ |
| *D. hydei* | 0.99 | 0.25 ± 0.03<sup>ab</sup> | -1.78 ± 0.18 | 1.03 ± 0.12 | 3.98 ± 0.41 | 327 ± 193 |
| *D. melanogaster* | 0.99 | 0.25 ± 0.02<sup>ab</sup> | -0.03 ± 0.14 | 0.40 ± 0.05 | 4.04 ± 0.39 | 298 ± 163 |
| *D. sulfurigaster* | 0.94 | 0.19 ± 0.02<sup>b</sup> | 1.79 ± 0.54 | 0.37 ± 0.14 | 5.25 ± 0.54 | 80 ± 36 |
| *D. teissieri* | 0.98 | 0.17 ± 0.02<sup>b</sup> | 3.60 ± 0.38 | 0.25 ± 0.07 | 5.75 ± 0.67 | 54 ± 25 |

\*The accompanying standard error is calculated using the delta method and does not represent measured variation.

Slopes of the TDT curves were used to calculate Q_10_’ values for how injury accumulation rate increases with decreasing temperature. Here we found Q_10_’s ranging from 54 to 6454 (**Table 2**), highlighting that even small changes in exposure temperature affect the injury accumulation rate dramatically and by extension changes the duration that the cold stress can be tolerated substantially.

To examine if and how the TDT parameters of thermal sensitivity varied with the environmental conditions obtained from the species’ natural distribution range, we correlated TDT slopes against the average temperature of the coldest quarter and the average latitude across species’ habitats (see **Table 1**). Spearman’s correlation tests revealed no significant relationship between TDT slopes and either average temperature of the coldest quarter (Spearman’s ρ = -−0.657, P = 0.175; **Fig. 4A**) or average latitude (Spearman’s ρ = 0.829, P = 0.058; **Fig. 4D**), suggesting that the observed differences in slope do not correlate strongly to species biogeography.

**Figure 4.**
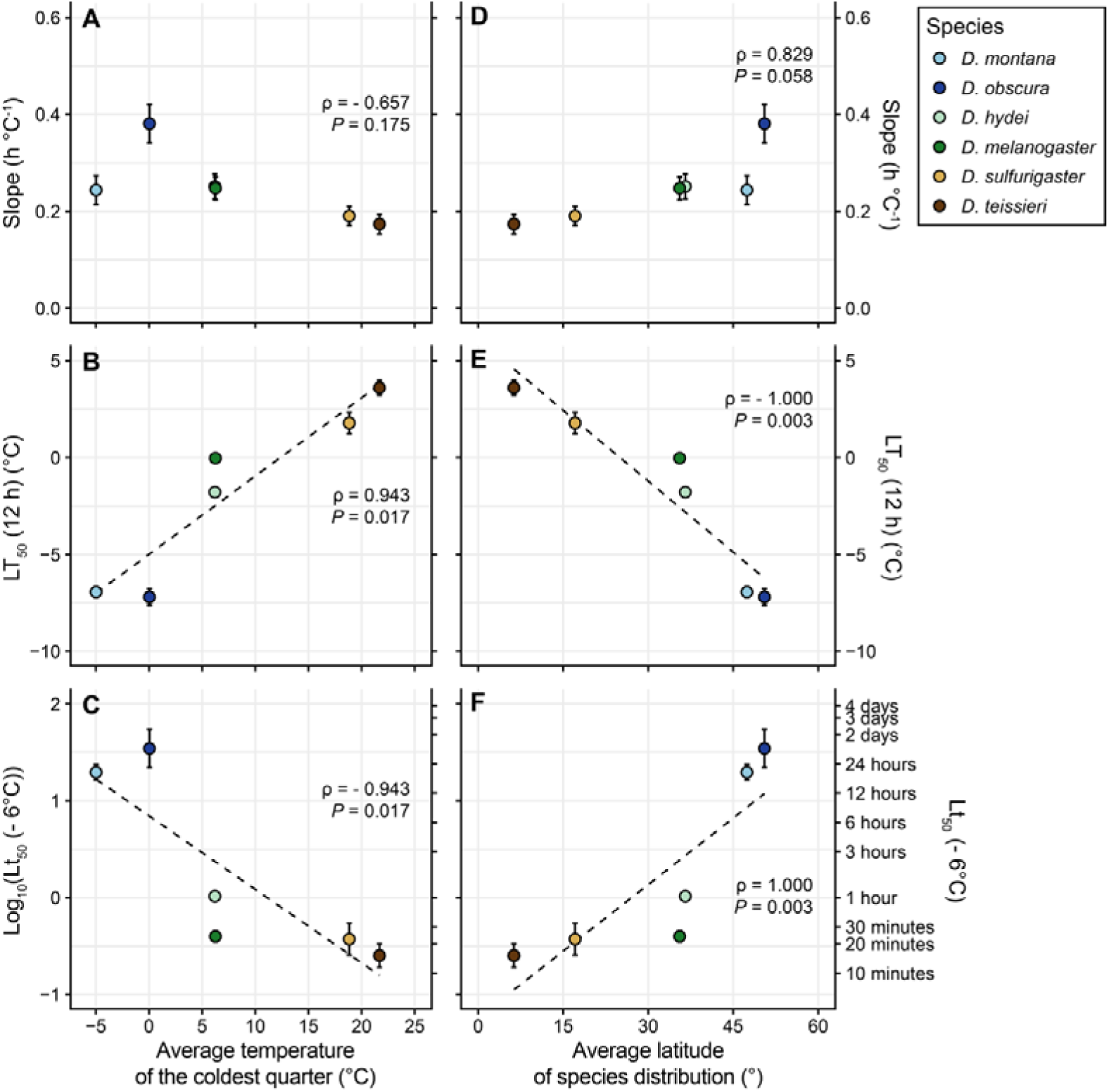
– Correlation analysis of thermal death time (TDT) parameters and species biogeography. Spearman’s correlation analysis between the average temperature of the coldest quarter for each species’ geographical distribution (left column, A-C), the average latitude for each species’ geographical distribution (right column, D-F), and species-specific parameters extracted from the TDT curves: The slope (A,D), LT50 (12h) (B,E), and Lt50 (−6°C) (C,F). Dashed lines represent potential linear regressions for statistically significant correlations, and error bars not visible are obscured by the symbols.

However, when examining the association between climatic or geographic variables with basal cold tolerance (i.e., the TDT intercepts at either −6 °C or 12 h; **Fig. 3**; **Table 2**) we found strong and significant correlations. LT_50_ (12 h) was strongly positively correlated with the average temperature of the coldest quarter (Spearman’s ρ = 0.943, P = 0.017; **Fig. 4B**) and strongly negatively correlated with latitude (Spearman’s ρ = - 1.000, P = 0.003; **Fig. 4E**). Similarly, Lt_50_ (at −6 °C) showed a strong negative correlation with the average temperature of the coldest quarter (Spearman’s ρ = –0.943, P = 0.017; **Fig. 4C**), while a strong positive correlation was found with the average latitude (Spearman’s ρ = 1.000, P = 0.003; **Fig. 4F**). Together, these results indicate that species from colder, high-latitude environments exhibit greater basal cold tolerance than species from warmer, low-latitude regions.

### Cold damage additivity

To test whether cold damage at different stressful temperatures is additive, flies were exposed to species-specific combinations of intense cold stress (T_1_ – the temperature causing LT_50_ in 30 min) and moderate cold stress (T_2_ – the temperature causing LT_50_ in 300 min) designed to cause 50% mortality (**Fig. 5**; **Table S2**). If cold stress-induced mortality was additive, we would expect no differences between groups, irrespective of the cold exposure combination.

**Figure 5.**
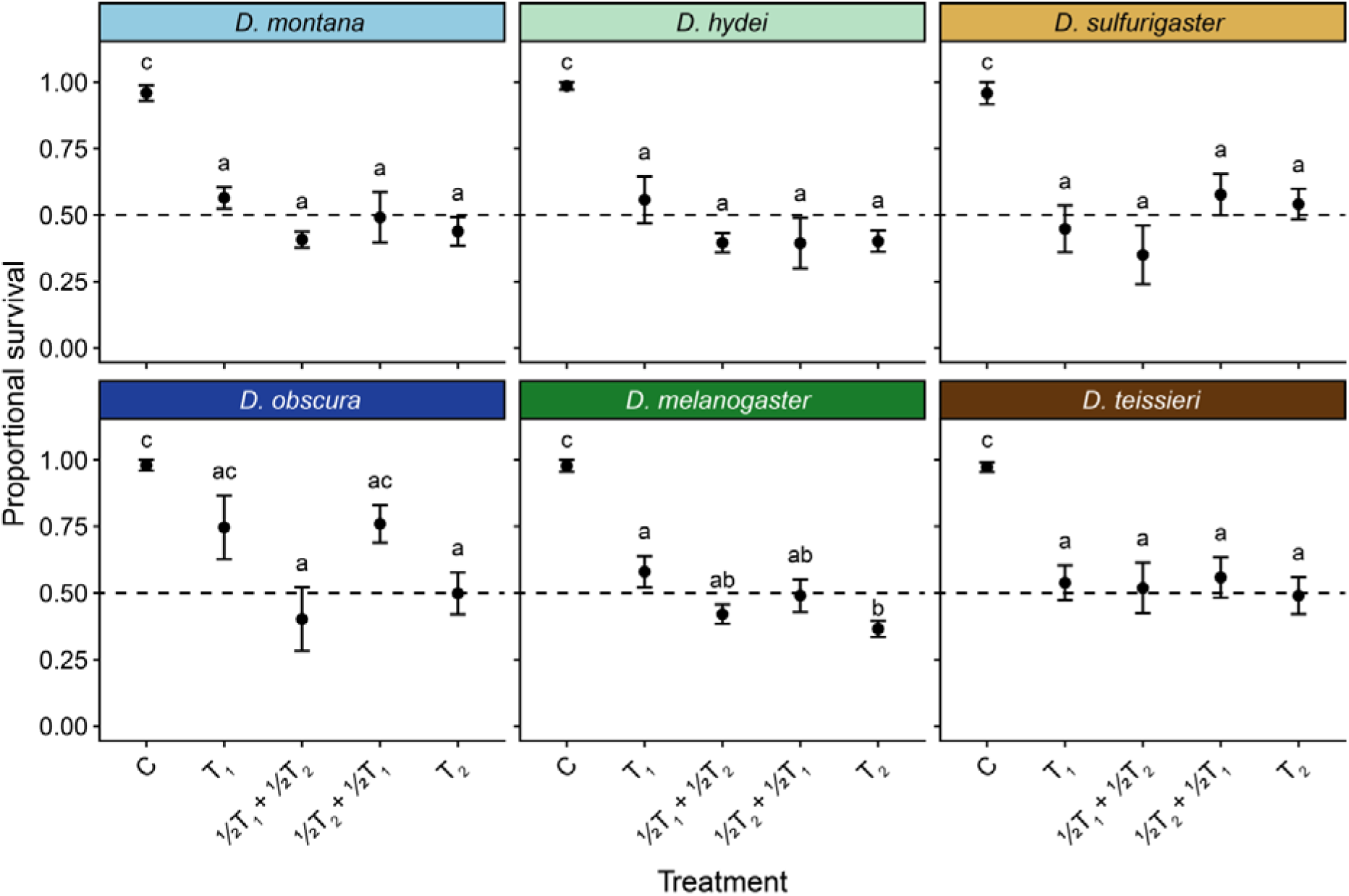
– Cold damage additivity. Average proportional survival after cold treatment for all six Drosophila species: T_1_ = Intense cold stress for 30 minutes, T_2_ = Moderate cold stress for 300 minutes, ½T_1_+½T_2_ = Intense cold stress for 15 minutes followed by moderate cold stress for 150 minutes, and ½T_2_ + ½T_1_ = Moderate cold stress for 150 minutes followed by intense cold stress for 15 minutes. Flies in the control group (C) were handled but received no cold treatment. Treatment groups sharing a letter (within each species) are not statistically different – note that lower case ‘c’ does not refer to the control group but is a letter indicating group differences. Temperature-specifics and sample sizes for each species and treatment group can be found in **Table S2**.

For all six species, fly mortality was scattered around 50% mortality for both the isolated T_1_ and T_2_ treatments and for the combinations, supporting the hypothesis that cold mortality at different stress intensities is additive (**Fig. 5**) and this was true regardless of the sequence of the two exposures (acute stress before or after chronic stress). In one species (*D. melanogaster*) the isolated exposure to T_1_ and T_2_ differed slightly but significantly; however mortality in the combined treatments for this species were intermediate to the two temperatures supporting the assumption of additivity. In another species (*D. obscura*) we noted that some of the treatment groups did not differ statistically from the controls, likely due to the large degree of variation within this species, but importantly the analysis revealed no differences between cold treatment groups, consistent with additivity.

### Longevity

To demonstrate that the relationship between survival duration and temperature is different in the non-stressful permissive temperature range we measured longevity at four different permissive temperatures, 11, 15, 19, and 25°C (**Fig. 6**). At permissive temperature fly mortality generally increased with increasing temperature such that life expectancy of the ectothermic flies decreases at warmer temperature as expected. Across species and temperatures we estimated Lt_50_ ranging from 10 to 166 days (**Fig. 6A**) and a TDT analysis within species find negative slopes (mean ± standard deviation across species = −0.049 ± 0.016 log_10_(Lt_50_) °C^-1^) corresponding to Q_10_ values for rate of mortality (1/time to death) ranging from 2.14 to 5.95 (**Fig. 6B**).

**Figure 6.**
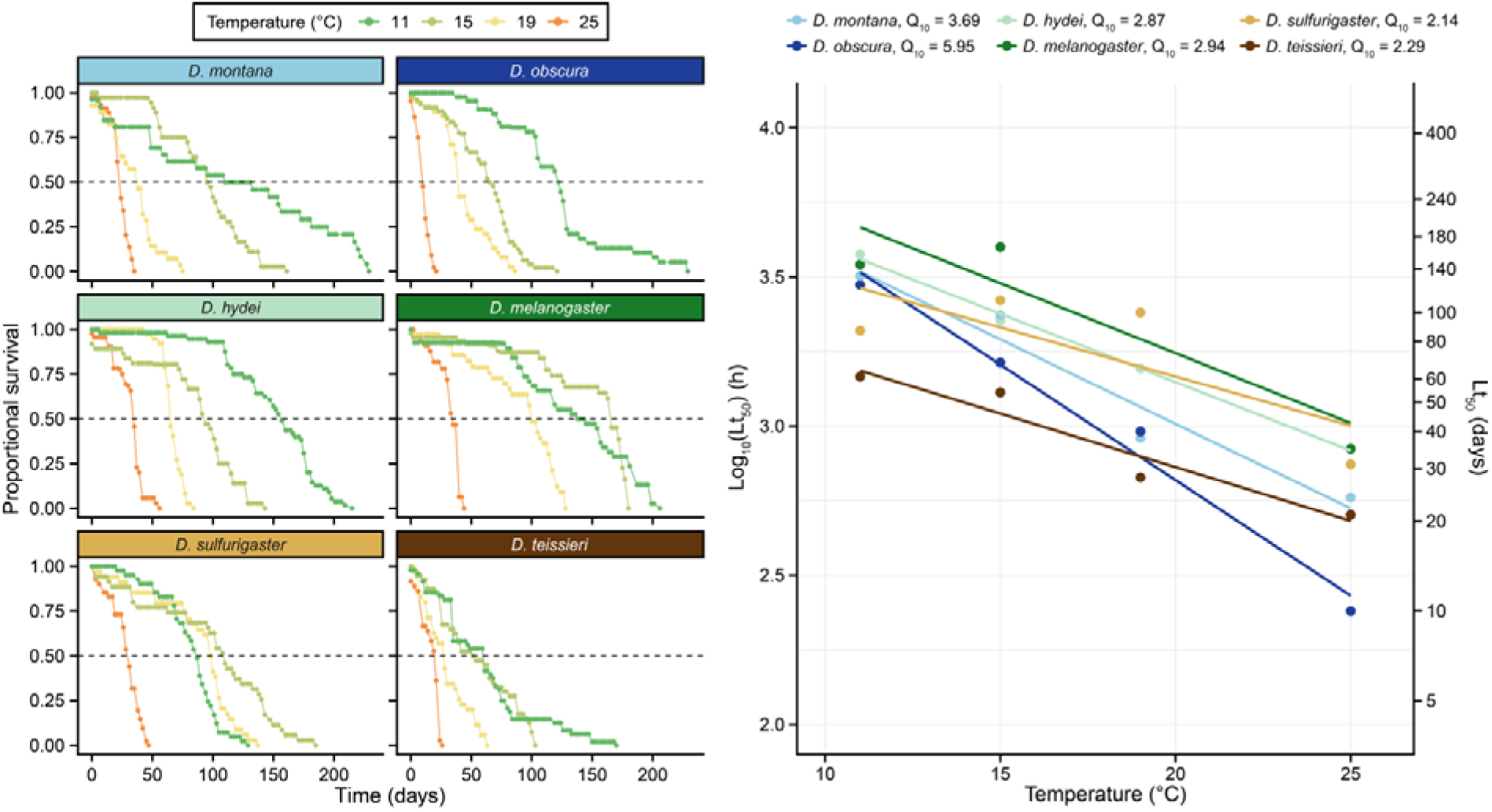
– Fly longevity and mortality at permissive temperatures. A) Proportional survival plotted against exposure time at 11, 15, 19, and 25°C for all six *Drosophila* species. The Lt_50_ was estimated as the day mortality surpassed 50% (i.e. the points directly below the dashed horizontal line). B) Log_10_ (Lt_50_) estimates plotted against exposure temperature with mean Q_10_ values for the rate of ageing for each species being calculated from the slope of the linear regression lines fitted to the data points as for the TDT analysis.

While the analysis revealed species-level differences in basal longevity (effect of species: F_5,12_ = 7.2, P = 0.003), the effect of temperature was similar across species (effect of species x temperature: F_5,12_ = 2.0, P = 0.148), supporting that the six species have similar thermal sensitivity of longevity within the permissive temperature range. Even so, we noted that the boreal species *D. obscura* appears challenged in regard to survival at 25°C, while the tropical species *D. sulfurigaster* and *D. teissieri* appear to find 11°C mildly stressful as indicated by their shorter Lt_50_’s at these temperatures (**Fig. 6B**).

## Discussion

Here we used TDT models to analyze cold tolerance of six *Drosophila* species originating from different climatic regions and thus exhibiting different levels of cold adaptation. The TDT data (Log_10_(Lt_50_) vs. temperature) from all 6 species were well fitted by the exponential model that provided a mathematical description of the relation between stress intensity (temperature) and tolerance duration (survival times ranging from ∼ 20 min to ∼2 days). Slopes of the TDT curves were similar for most species suggesting that the underlying physiological dysfunction associated with cold stress is similar across *Drosophila* species. In contrast, the intercepts of the TDT curves differed markedly among species with a clear horizontal shift in TDT curves that describe interspecific differences in cold resistance.

### Horizontal displacement of TDT curves

To evaluate differences in cold resistance we calculated where the TDT curve would intercept along the temperature (x-)axis (Rezende et al., 2014; Jørgensen et al., 2021). Specifically, we calculated the temperature that would cause Lt_50_ after 12 h from as a proxy for a limit representing the minimal temperature tolerated for a cold night (Fig. 3; **Table 2**). A similar analysis was made for the y-intercept, where we calculated how long species could survive a fixed temperature (−6 °C) (**Fig. 3**; **Table 2**). The parametric value for this latter calculation is, however, of limited ecological relevance for some species considering that tropical *Drosophila* will never experience such low temperature (e.g. mean temperature of the coldest quarter, see **Table 2**). Irrespective of analysis (x or y intercept) we found interspecific variance in cold tolerance to be tightly associated with the thermal environment from where the species originate (average temperature of the coldest quarter) and therefore also tightly linked to latitudinal distribution (**Fig. 4**). These strong correlations are entirely consistent with similar correlative investigations in large comparative insect studies (Addo-Bediako et al., 2000; Sunday et al., 2011; Sunday et al., 2019) including studies focused on *Drosophila* specifically (Ohtsu et al., 1998; Kimura, 2004; Strachan et al., 2011; Kellermann et al., 2012a; Andersen et al., 2015; Moos et al., 2025).

In quantitative terms we find the slope of Lt_50, 12h_ against average temperature of the coldest quarter to be ∼ 0.4 °C °C^-1^ such that basal cold tolerance increases 0.4 °C for every degree colder the species-specific environment is. Such slopes are also found using much larger compilations of *Drosophila* species (see **Fig. 7**) (Ohtsu et al., 1998; Kimura, 2004; Kellermann et al., 2012a; Andersen et al., 2015; MacLean et al., 2019; Moos et al., 2025). In these large-scale studies the lethal cold tolerance limits generally increases by 0.4 – 0.5 °C for every degree colder the winter temperature at the species-specific origins (mean = 0.45 °C °C^-1^), while the functional cold limit to animal performance (i.e. the CT_min_) has a somewhat lower slope (mean = 0.27 °C °C^-1^).

**Figure 7.**
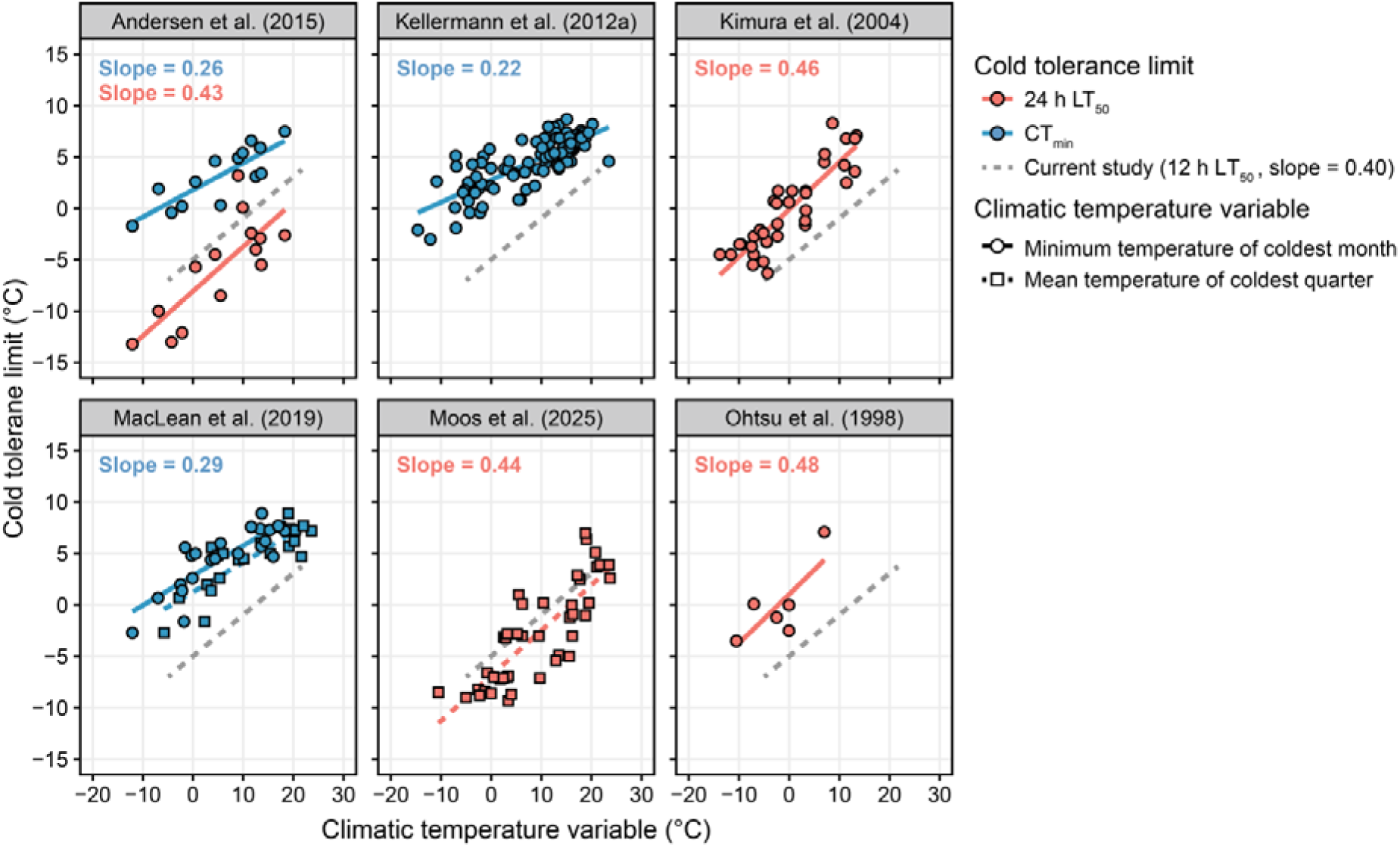
– Comparison of species-specific climate variables and cold tolerance limits across *Drosophila* studies. A comparative analysis reveals that the relationship between the degrees of cold experienced by *Drosophila* species in their natural the environments and cold tolerance is not only strong but highly similar across studies. Notably, the slopes of the relationship are shallower for the functional limit to animal performance (CT_min_, blue) than it is for lethal limits (LT_50_, red) regardless of whether the climate variable is the lowest experienced temperature across the year (circles, filled line) or a measure of seasonal cold (squares, dashed line). Data from the current study (12 h LT_50_) is indicated by the dashed grey line (slope = 0.40). Climatic temperature variables for studies where these were not reported were obtained using the same approach as Moos et al. (2025). Note that these slopes have not been phylogenetically corrected.

The present study, as well as studies highlighted in **Fig. 7**, use species that are reared at identical thermal conditions such that the observed interspecific differences in cold tolerance do not consider how species from boreal and temperate regions would naturally also be acclimated or acclimatized to lower temperatures in winter. Comparative examinations of the cold acclimation response, using compilations of tropical, temperate and boreal *Drosophila,* find critical limits (CT_min_) to decrease by 0.3 to 0.5 °C per °C of lowered acclimation temperature (Sørensen et al., 2016; Schou et al., 2017) and cold acclimation has directly been shown to horizontally displace TDT curves for cold tolerance in *D. suzukii* (Tarapacki et al., 2021). Interestingly, this level of plasticity of *Drosophila* is higher than that reported more generally for terrestrial ectotherms and insects in general (Gunderson and Stillman, 2015; Weaving et al., 2022). Nevertheless, combining the observed interspecific differences in *Drosophila* cold tolerance (∼ 0.45 °C per °C lower temperature of the environmental origin) with the plastic effects expected from seasonal cold acclimation (∼ 0.4 °C per °C lower acclimation temperature in the environment, see Schou et al. (2017) and MacLean et al. (2019)) would explain almost perfectly how boreal species are able to survive lower winter temperature, as the additive effects of adaptation and acclimation would explain ∼ 0.85 °C displacement in cold resistance per °C colder environment. An emergent observation from this rough and somewhat naïve analysis would then suggest that *Drosophila* species, across tropical to boreal environments, are adapted and acclimated to endure similar degrees of cold stress when considering their natural thermal habitat. This observation would therefore imply that species may be distributed as far towards high latitudes and cold climates as their cold tolerance allows. Several complimentary observations support this general suggestion. Firstly, as also observed in the present study, the association between tolerance and distribution is remarkably strong when investigated across *Drosophila* species (**Fig. 7**). Second, insect species in general (not *Drosophila* specifically) have been found to change distributional limits towards higher latitudes and elevation in close association with the increase in environmental temperature caused by global warming (Parmesan et al., 1999; Sunday et al., 2012; Freeman et al., 2018). Finally, cold tolerance will intuitively play a larger role for geographical persistence at the low end of the temperature spectrum since the potential for dynamic behavioral avoidance is often very limited at low temperature. This is somewhat different from the situations at high temperature extremes where behavioral avoidance is likely highly implicated in adaptations that determine distributional patterns (Kearney et al., 2009; Sunday et al., 2014). In accordance, the association between heat tolerance limits (CT_max_) and temperature of the environment are generally weaker (see for example Kellermann et al. (2012b), Addo-Bediako et al. (2000), and Kimura (2004)) than the strong association between cold tolerance and environment.

### Similarity of TDT slopes across species – same stress, different temperature

The slope of a TDT model for cold tolerance depicts the factorial decrease in tolerance duration (Lt_50_) with decreasing temperature. Because Lt_50_ is inversely related to the rate of injury accumulation, the slope effectively describes how the rate of cold injury increases exponentially with decreasing temperature. In TDT literature the slope is often expressed as a z value (or z’ for cold) where z’ = 1/slope, signifies how many degrees temperature should change along the TDT curve for a 10-fold change in survival time (Rezende et al., 2014; Jørgensen et al., 2021). Both the slope or z’ therefore express thermal sensitivity of the biological rates underlying cold injury and they are therefore homologous to Q_10_ (Q_10_’ for cold – see methods). Classical Q_10_ analysis reports the *increase* in biological rates with increasing temperature while the Q_10_’ parametrizes the *decrease* in injury rate that occurs with increasing temperature. This is consistent with a physiological interpretation where the increase in cold injury rate is caused by the loss of a homeostatic biological rate such that injury rate increases when the homeostatic rate decreases and *vice versa* (see discussion below)

Here we find steep slopes (**Fig. 3**) and therefore high Q_10_’ values (**Table 2**) for all species. One boreal species (*D. obscura*) had a significantly higher thermal sensitivity (slope) than the two tropical species (*D. sulfurigaster and D. teissieri*), but there was no systematic significant association between either the thermal (**Fig. 4A**) or latitudinal (**Fig. 4D**) parameters that characterize the natural habitat of the species examined. Thermal sensitivity of cold injury rate was characterized by Q_10_’ ranging from 54 to 327 in the five more similar species and Q_10_’ was estimated at 6454 for the outlier species *D. obscura* (**Table 2**). For all six species these values demonstrate that cold injury rate has a high thermal sensitivity which is consistent with previous estimates from *Drosophila* species. For example, Q_10_’ values calculated from z’ ranged from 31 to 175 in *D. suzukii* acclimated to different temperatures (Tarapacki et al., 2021) and a Q_10_’ of ∼70 can be calculated for *D. melanogaster* based on data from (Chen and Walker, 1994). Further, Rezende et al. (2014) reported z’ values for cold injury in other insect taxa that were also generally high such that the calculated Q_10_’ span from ∼ 2 in *Sitophilus granarius* (Fields, 1992) to ∼ 46 in *Stegobium paniceum* (Abdelghany et al., 2010).

Considering that Q_10_ for “normal” biological rates typically range from 1.5–3 (Angilletta, 2009; Cossins, 2012; Seebacher et al., 2015; Jørgensen et al., 2022), the Q_10_’ for chill injury rate in insects has an atypical high thermal sensitivity. To contrast the thermal sensitivity of cold stress to biological rates at permissive temperatures we measured longevity (similar to Lt_50_) in the temperature range between 11–25 °C (**Fig. 6**). In this range the slopes of the TDT curves became negative, and the Q_10_ values that represent the rate of senescence/ageing (Q_10_ ranging from 2.14-5.95) were closer to those normally reported for biological rates of ectotherms (**Fig. 6**; see Dell et al. (2011) and Ørsted et al. (2022)). Accordingly, our TDT curves constructed at low temperatures causing cold stress (**Fig. 2**) provides reliable model estimates of how cold injury accumulation intensifies with decreasing temperatures, but as seen at more permissive temperature, these models have no resemblance to the factors shaping longevity at permissive temperature. We therefore caution that TDT model parameters are not extrapolated to irrelevant thermal ranges where thermal stress ceases to be the primary cause of mortality (see discussion in Jørgensen et al. (2021) and Ørsted et al. (2022)). From an ecological perspective the high Q_10_’ values related to chill injury rate signify that insects may operate at narrow thermal margins with respect to cold stress and that modest decreases in temperature will markedly accelerate injury rate. As discussed above, this is consistent with the observation that increased winter temperatures associated with global warming has allowed for distributional shifts towards higher latitudes or elevations (Parmesan et al., 1999; Sunday et al., 2012; Freeman et al., 2018).

From a physiological perspective, the atypical high Q_10_’ related to cold injury rate suggests that the underlying physiological rates are also atypical. Since all species examined here were characterized by high Q_10_’ this suggest that that there is a commonality in their cold stress “syndrome” even if the stress caused by cold occurs at different temperature ranges for tropical and boreal species. In fact, the analysis of slope/thermal sensitivity bears some resemblance to an Arrhenius-type analysis where breakpoints in a TDT curve would indicate shifts in thermal sensitivity that imply different underlying physiological processes (Cossins, 2012; Somero et al., 2017), while continuous linearity, as found here, is supportive of a common physiological mechanism across the range of temperatures examined (although this is not concrete proof). To further examine if the same physiological processes underlie failure at both high and moderate cold stress intensities we examined if stress at different intensities is additive (**Fig. 5**) (Fry et al., 1946; Jørgensen et al., 2021). Our results support the additive model for all species with additive injury accumulation from moderate and intense stress (**Fig. 5**) supporting the hypothesis that cold-induced mortality across the stressful range arises from shared physiological mechanisms.

The commonality in phenotype between species that were all characterized by high Q_10_’ and additivity of cold stress, suggest that similar physiological dysfunctions could be the cause cold mortality regardless if it observed at, for example 8 °C for tropical *D. teissieri* or at –13 °C for boreal *D. montana* (**Fig. 3**). The present study has no concrete data to support the idea of a common cold stress syndrome, but numerous studies, also involving *Drosophila*, have demonstrated that cold-induced injury in insects is associated with a gradually developing homeostatic collapse (MacMillan et al., 2015b; MacMillan, 2019; Overgaard et al., 2021). Specifically, an impaired ability to maintain ion and water homeostasis in cold-exposed insects causes neuromuscular dysfunction and ultimately mortality. This common cold stress syndrome is characterized by a progressive rise in hemolymph KL caused by NaL leakage into the gut with osmotically-following water (Koštál et al., 2004; MacMillan et al., 2012; Lebenzon et al., 2020), which lowers hemolymph volume and increases KL concentration (MacMillan et al., 2015b). This is further exacerbated by passive K^+^ diffusion into the hemolymph due to reduced active transport capacity (Andersen et al., 2017; Andersen and Overgaard, 2020). The resulting extracellular hyperkalemia causes depolarization of excitable cell membranes, trapping insects in a state of chill coma and, if prolonged, contributing to chilling injury and subsequent mortality (reviewed in Overgaard et al. (2021)). Whether this cold stress syndrome shows a similar progression across different, stressful temperatures and across species remains to be explored systematically and in detail.

In conclusion our results show that TDT models capture cold-tolerance differences among *Drosophila* species through clear shifts in tolerance thresholds rather than changes in thermal sensitivity. The similarity in TDT slopes support a common physiological mechanism underlying cold injury in these chill susceptible insects, while the consistent interspecific differences in intercepts highlight how evolution of cold tolerance has shifted the physiological tolerance such that cold adapted species only experience the same degree of stress once they are exposed to a lower temperature. TDT analysis represents a laborious experimental approach to study cold biology, but as discussed here this approach allows for both physiological and ecological inference. We note however, that simple assays to estimate thermal tolerance (e.g. CT_min_) captured very similar patterns of interspecific variance in cold tolerance and we argue that the use of TDT in evolutionary and ecological studies should be evaluated while considering the cost/benefits of this rigorous and detailed analysis.

## Supplementary material

**Table S1.**
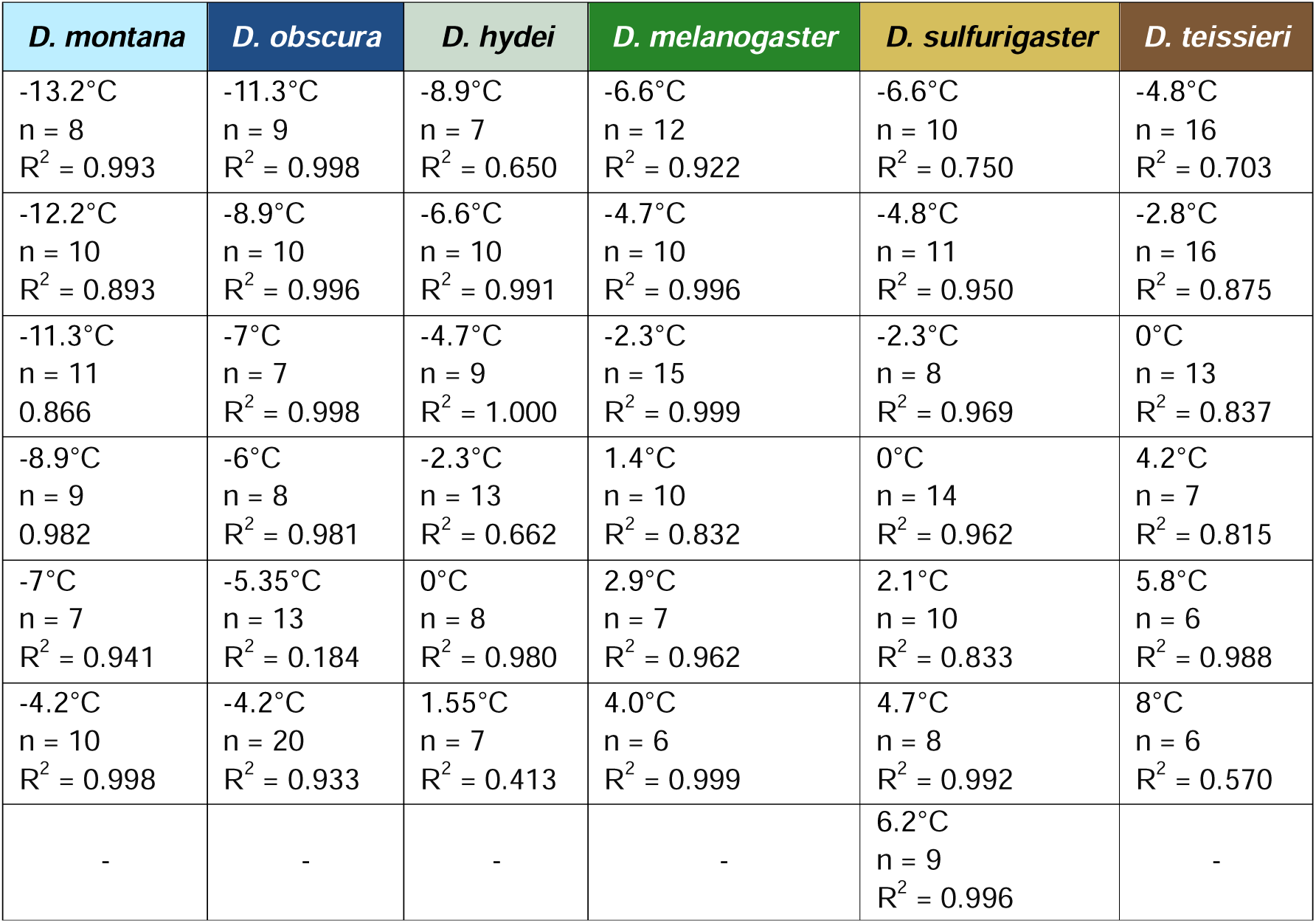
– Experimental temperature, number of replicates, and R^2^ explaining the fit of the Lt_50_ data points to the logistic regression model for the six *Drosophila* species.

| *D. montana* | *D. obscura* | *D. hydei* | *D. melanogaster* | *D. sulfurigaster* | *D. teissieri* |
| --- | --- | --- | --- | --- | --- |
| -13.2°C<br>n = 8<br>$R^2 = 0.993$ | -11.3°C<br>n = 9<br>$R^2 = 0.998$ | -8.9°C<br>n = 7<br>$R^2 = 0.650$ | -6.6°C<br>n = 12<br>$R^2 = 0.922$ | -6.6°C<br>n = 10<br>$R^2 = 0.750$ | -4.8°C<br>n = 16<br>$R^2 = 0.703$ |
| -12.2°C<br>n = 10<br>$R^2 = 0.893$ | -8.9°C<br>n = 10<br>$R^2 = 0.996$ | -6.6°C<br>n = 10<br>$R^2 = 0.991$ | -4.7°C<br>n = 10<br>$R^2 = 0.996$ | -4.8°C<br>n = 11<br>$R^2 = 0.950$ | -2.8°C<br>n = 16<br>$R^2 = 0.875$ |
| -11.3°C<br>n = 11<br>0.866 | -7°C<br>n = 7<br>$R^2 = 0.998$ | -4.7°C<br>n = 9<br>$R^2 = 1.000$ | -2.3°C<br>n = 15<br>$R^2 = 0.999$ | -2.3°C<br>n = 8<br>$R^2 = 0.969$ | 0°C<br>n = 13<br>$R^2 = 0.837$ |
| -8.9°C<br>n = 9<br>0.982 | -6°C<br>n = 8<br>$R^2 = 0.981$ | -2.3°C<br>n = 13<br>$R^2 = 0.662$ | 1.4°C<br>n = 10<br>$R^2 = 0.832$ | 0°C<br>n = 14<br>$R^2 = 0.962$ | 4.2°C<br>n = 7<br>$R^2 = 0.815$ |
| -7°C<br>n = 7<br>$R^2 = 0.941$ | -5.35°C<br>n = 13<br>$R^2 = 0.184$ | 0°C<br>n = 8<br>$R^2 = 0.980$ | 2.9°C<br>n = 7<br>$R^2 = 0.962$ | 2.1°C<br>n = 10<br>$R^2 = 0.833$ | 5.8°C<br>n = 6<br>$R^2 = 0.988$ |
| -4.2°C<br>n = 10<br>$R^2 = 0.998$ | -4.2°C<br>n = 20<br>$R^2 = 0.933$ | 1.55°C<br>n = 7<br>$R^2 = 0.413$ | 4.0°C<br>n = 6<br>$R^2 = 0.999$ | 4.7°C<br>n = 8<br>$R^2 = 0.992$ | 8°C<br>n = 6<br>$R^2 = 0.570$ |
| - | - | - | - | 6.2°C<br>n = 9<br>$R^2 = 0.996$ | - |

**Table S2.**
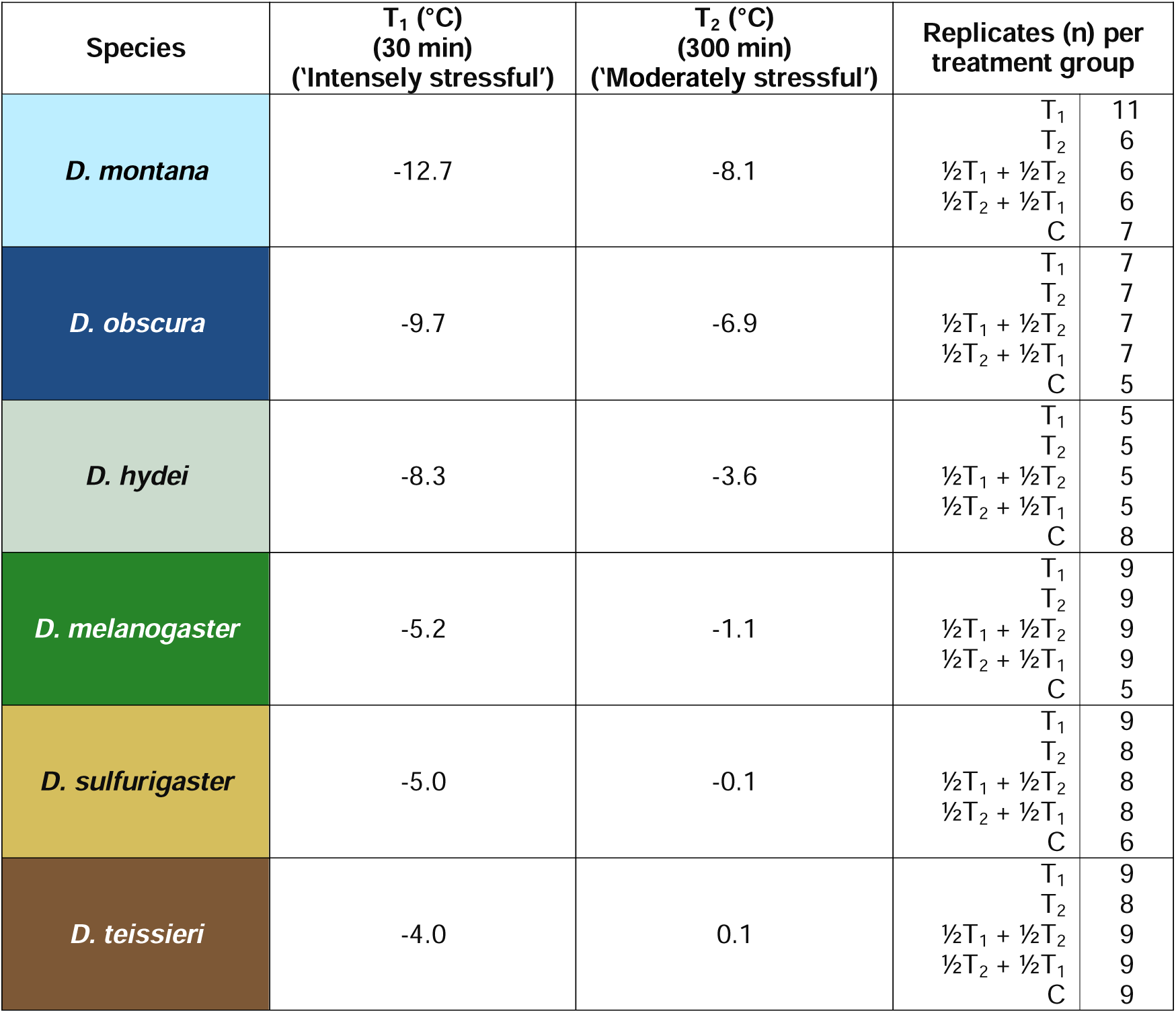
- Experimental temperatures for the two types of cold stress provided in the additivity study and the number of replicates per treatment group for the six *Drosophila* species.

